# The symbiotic complex of *Dendroctonus simplex*: implications in the beetle attack and its life cycle

**DOI:** 10.1101/276857

**Authors:** Audrey-Anne Durand, Philippe Constant, Eric Déziel, Claude Guertin

## Abstract

The eastern larch beetle (*Dendroctonus simplex* Le Conte) is recognized as a serious destructive forest pest in the upper part of North America. Under epidemic conditions, this beetle can attack healthy trees, causing severe damages to larch stands. *Dendroctonus* species are considered as holobionts, as they engage in multipartite interactions with microorganisms, such as bacteria, filamentous fungi, and yeasts, which are implicated in physiological processes of the insect, such as nutrition. They also play a key role in the beetle’s attack, as they are responsible for the detoxification of the subcortical environment and weaken the tree’s defense mechanisms. The eastern larch beetle is associated with bacteria and fungi, but their implication in the success of the beetle remains unknown. Here, we investigated the bacterial and fungal microbiota of this beetle pest throughout its ontogeny (pioneer adults, larvae and pupae) by high-throughput sequencing. A successional microbial assemblage was identified throughout the beetle developmental stages, reflecting the beetle’s requirements. These results indicate that a symbiotic association between the eastern larch beetle and some of these microorganisms takes place and that this *D. simplex* symbiotic complex is helping the insect to colonize its host tree and survive the conditions encountered.

## Introduction

The eastern larch beetle, *Dendroctonus simplex* LeConte (Coleoptera: Scolytinae), is a phloem-feeding insect that attacks tamarack trees, *Larix laricina* (Du Roi) K. Koch (Wood, 2007). During the dispersal period, pioneer beetles attack trees and build galleries in the phloem layer. Following the reproduction and eggs hatching, larvae also excavate galleries and eat phloem throughout their development. The last larval instar digs a pupal chamber, stops feeding and empties his digestive tract in preparation for transformation in pupae, representing an inactive stage. Pupae will than transform into adults, overwintering until the next dispersal period (Langor & Raske, 1987a; Langor & Raske, 1987b). This beetle is considered a secondary pest because it usually attacks weakened or freshly dead trees. However, under epidemic conditions, it can attack healthy trees, causing severe damage to larch stands (Langor & Raske, 1987a; Langor & Raske, 1989b). The widespread attacks of the eastern larch beetle observed in the past suggest that this pest should no longer be considered secondary (Langor & Raske, 1989a). More recently, extensive attacks have been observed in Quebec (Canada), causing the death of thousands of larch trees across the province.

Bark beetles engage in a multitude of interactions with various microorganisms to form a holobiont (Margulis & Fester, 1991; Six, 2013). *Dendroctonus* species are commonly associated with bacteria, filamentous fungi, and yeasts. This assemblage of microorganisms, also called microbiota, is implicated in various physiological processes of the insect, colonization of the host tree, and protection from antagonistic organisms (Popa *et al.*, 2012; Hofstetter *et al.*, 2015). Accordingly, microorganisms included in the *Dendroctonus* microbiota may act as key factors in their success.

*Dendroctonus* species are associated with a variety of bacterial genera. Some of these bacteria are implicated in the beetle nutrition, supplementing the nutrient-poor phloem diet with amino acids, nitrogen and vitamins (Bridges, 1981; Morales-Jimenez *et al.*, 2009; Morales-Jimenez *et al.*, 2013). Some are also able to break down cellulose, helping the assimilation by the insect (Morales-Jimenez *et al.*, 2012). Additionally, some bacteria are implicated in detoxification processes, such as terpenoid degradation (Adams *et al.*, 2013). Some bacteria also produce antifungal compounds that protect the insect against antagonistic fungi (Scott *et al.*, 2008). On the other hand, associated filamentous fungi are also implicated in beetle nutrition, supplementing their diet with sterols (Bentz & Six, 2006). Some fungi can weaken the tree defense system, leading the insect to a successful attack (Paine *et al.*, 1997). Yeasts also play a significant role in beetle success, as they are reported to be implicated in nutrition, detoxification of plant defense compounds and protection from antagonistic microorganisms (Davis, 2014). Additionally, associated yeasts are referred as mediators of the beetle-microorganisms interactions (Davis *et al.*, 2011; Davis, 2014; Hofstetter *et al.*, 2015). It appears that several functions are redundant among microorganisms forming the microbiota.

In the present study, the bacterial and fungal microbiota of the eastern larch beetle was identified at various beetle developmental stages (pioneers adult, larvae, and pupae) by high-throughput sequencing of the 16S and 18S rRNA genes. Since they are distinct, as we have previously demonstrated, both the ecto- and the endomicrobiota of the insect were investigated (Durand *et al.*, 2015). The aim of this study was to understand the crucial role played by the microbiota of *D. simplex* in the colonization success and to follow the changes that may occur in the microbial populations throughout its life cycle. We hypothesized that a core microbiota should be associated with *D. simplex* and that the proportion of these microorganisms should vary in function of beetle development. Moreover, pioneer beetles should carry microorganisms playing a key role in the beetle attack. These results will help us understand the eastern larch beetle attack and colonization process.

## Materials and Methods

### Site location, beetles processing, and samples preparation

Insects were collected from a Quebec Province larch plantation located near Saint-Claude (Quebec, Canada; Lat. 45.6809, Long. −71.9969) with the permission of the *Ministère des Forêts, de la Faune et des Parcs* authority. Log sections of randomly selected larch trees showing apparent signs of attacks by *D. simplex* were transported to the laboratory where they were stored at room temperature in plexiglass cages (30 cm ×30 cm ×88 cm). Beetle development monitoring and retrieval were achieved as described previously (Durand *et al.*, 2017). For each developmental stage, microorganisms associated with the ecto- and endomicrobiota were separated and recovered as previously described (Durand *et al.*, 2015). For the ectomicrobiota of each development stage, 50 insects per replicate were randomly selected, and pooled in a 15 ml polypropylene tube to recover sufficient bacterial genomic DNA from the surface of the cuticle. Then, each sample underwent five successive washes with 5 ml phosphate-buffered saline (PBS) containing 0.1 % Triton X-100, with 1 min agitation (Genie 2 Vortex, Fisher, Ottawa, ON, Canada). The solution was filtered through a 0.22 μm nitrocellulose filter (EMD Millipore, Billerica, MA, USA) to recover the biomass.Each filter was placed in a Lysing matrix A tube (MP Biomedicals, Solon, OH, USA) for DNA extraction. Ten previously washed beetles were randomly selected for each replicate to retrieve the endomicrobiota. Their external surface was sterilized with three serial washing in 70% EtOH, followed by one wash with sterile ultrapure water, as this protocol as shown a complete separation of both microbiota before (Durand *et al.*, 2015). The insects were then crushed into PBS and placed in a 2 ml screw cap tube containing 200 mg of 0.1 mm glass beads (BioSpecs, Bartlesville, OK, USA) for DNA extraction. Three biological replicates were achieved for each developmental stage and microbiota, representing a total of 18 samples.

### DNA Extraction and PCR amplification

Total DNA was extracted following the method previously described (Durand *et al.*, 2015). A negative control using all the extraction solutions but no insect was achieved. DNA concentration was estimated using the Quant-iT(tm) PicoGreen* dsDNA Assay Kit (Invitrogen, Life Technologies, Burlington, ON, Canada) following the manufacturer instruction. The integrity of the genomic DNA was confirmed on a 1% agarose gel stained with ethidium bromide and visualized under UV light.

PCR amplification was achieved to confirm the presence of microbial DNA in each sample. For the bacteria, universal primers 27F (5’ AGA GTT TGA TCC TGG CTA G 3’) and 1492R (5’ GGT TAC CTT GTT ACG ACT T 3’) were used to amplify the 16S rRNA gene (Kwong & Moran, 2013). For the fungi, universal primer NSA3 (5’ AAA CTC TGT CGT GCT GGG GAT A 3’) and NLC2 (5’ GAG CTG CAT TCC CAA ACA ACT C 3’) were used to amplified the SSU, ITS, and LSU regions of the rRNA genes (Martin & Rygiewicz, 2005). Each 50 μl PCR reaction contained 25 mM MgCl2, 10 ^®^g BSA, 10 mM dNTPs, 10 mM of each primer, 5 U Taq DNA polymerase and ThermoPol^®^ buffer (New England Biolabs, Whitby, ON, Canada). For the bacteria, following the initial denaturation step of 5 min at 94°C, 30 amplification cycles were performed (94°C for 30 s, 55°C for 30 s, 72°C for 1 min and 30 s) followed by a final extension step at 72°C for 10 min. For the fungi, following the initial denaturation step of 5 min at 94°C, 30 amplification cycles were performed (94°C for 30 s, 67°C for 30 s, 72°C for 1 min) followed by a final extension step at 72°C for 10 min. Amplification was confirmed by electrophoresis of the PCR products on a 1.5% agarose gel stained with ethidium bromide and visualized under UV light. Bacterial and fungal DNA was present in each sample, except for the negative control.

### Pyrosequencing of microbial DNA

Genomic DNA from each sample was sent to Research and Testing Laboratory (Lubbock, TX, USA) for sequencing. The bacterial 16S rRNA gene was amplified using the universal primers 28F (5’ GAG TTT GAT CNT GGC TAC G 3’) and 519R (5’ GTN TTA CNG CGG CKG CTG 3’) targeting the V1-V3 hypervariable regions. Roche 454 FLX-Titanium chemistry was used to sequence the amplicons. Elongation was performed from the forward primer. Raw data are available on NCBI under BioProject number PRJNA401528.

Sequences related to fungal microbiota associated with the eastern larch beetle (BioProject PRJNA354793) presented in our previous study (Durand *et al.*, 2017) were also used in this study. The fungal 18S rRNA gene was amplified using the universal primers SSUForward (5’ TGG AGG GCA AGT CTG GTG 3’) and funTitSsuRev (5’ TCG GCA TAG TTT ATG GTT AAG 3’). Roche 454 FLX-Titanium chemistry was also used to sequence the amplicons. Elongation was performed from the forward primer as well.

### Sequences processing pipeline

The post-sequencing processing was completed using the open-source program mothur v.1.33.0 software (http://www.mothur.org) (Schloss *et al.*, 2009). For the sequences associated with the bacterial microbiota, the pipeline described by Comeau and collaborators was followed (Comeau *et al.*, 2012). Raw 454 reads were first processed to remove low-quality reads, such as (i) the presence of one or more uncertain bases (N), (ii) sequences shorter than 150 nt (nucleotides), (iii) unusually long reads that extended more than 100 nt over the amplicon size, (iv) reads that have long homopolymer sequences (more than 8), and (v) reads with incorrect forward primer sequences. Regions corresponding to the forward primer were kept to facilitate the alignment of the sequences during subsequent analyses. Contaminant sequences, such as chloroplast and mitochondria, were removed from the dataset. Additionally, chimeras were removed with UCHIME (Edgar *et al.*, 2011), as implemented in mothur. The remaining filtered sequences were aligned by domain against the SILVA reference alignment release 119 (Quast *et al.*, 2013) using the ksize=9 parameter in mothur. Reads were also trimmed of all bases beyond the reverse primer with BioEdit 7.2.5 (http://www.mbio.ncsu.edu/bioedit/bioedit.html). Singletons were finally removed after clustering into draft Operational Taxonomic Units (OTUs) to obtain the final quality reads. Libraries were normalized to the sequencing effort of the smallest 16S rRNA gene library (1744 sequences/samples) to avoid biases in comparative analyses introduced by the sampling depth. The last aligned reads were clustered into OTUs at ≥ 97% identity threshold using the furthest neighboring cluster in mothur (Schmitt *et al.*, 2012). Representative sequences of each OTU were taxonomically identified using the Ribosomal Database Project (RDP) classifier (Wang *et al.*, 2007).

For the sequences associated with the fungal microbiota, raw sequences processing, clusterization, taxonomical identification and equalization of the library was done as described before (Durand *et al.*, 2017).

### Data analysis

Rarefaction curves were generated within the mothur software to evaluate the sufficiency of the sequencing effort (data not shown). Shannon diversity index was also calculated with mothur, and ANOVA was performed with JMP Pro 12 (SAS Institute Inc., Cary, NC, USA) on obtained values. A PERMANOVA analysis on the generated bacterial and fungal OTUs was performed with the R software 3.1.3 (http://www.r-project.org) using the package “vegan”. A Euclidean distance matrix was used to generate a UPGMA agglomerative clustering according to the identified OTUs and their abundance with R using the package “hclust”. Significant nodes were identified by performing 999 permutations of the dataset. The heatmap representing OTUs abundance was generated with the package “gplots” and “RColorBrewer”.

## Results

### Diversity of the associated microbial community across *D. simplex* developmental stages

The microbial diversity associated with the eastern larch beetle developmental stages were compared using 18 samples that were sequenced to characterize the bacterial and fungal microbiota. For the bacteria, a total of 31,392 high-quality-filtered sequences were recovered after quality control and equalization. The average read length was 442 bp. Clusterization at a 97% pairwise-identity threshold generated 4009 OTUs. The Shannon diversity index was calculated for all samples, and significant differences (ANOVA test; *F* = 9.52; *p* < 0,0007) are observed throughout the beetle developmental stages (Table 1). The endomicrobiota of the larvae exhibited the lowest diversity, followed by the ectomicrobiota of all developmental stages. The highest diversity was associated with the pupae endomicrobiota. Overall, the Shannon diversity index is higher for the endomicrobiota than the ectomicrobiota, except for the larvae samples.

**Table 1.**
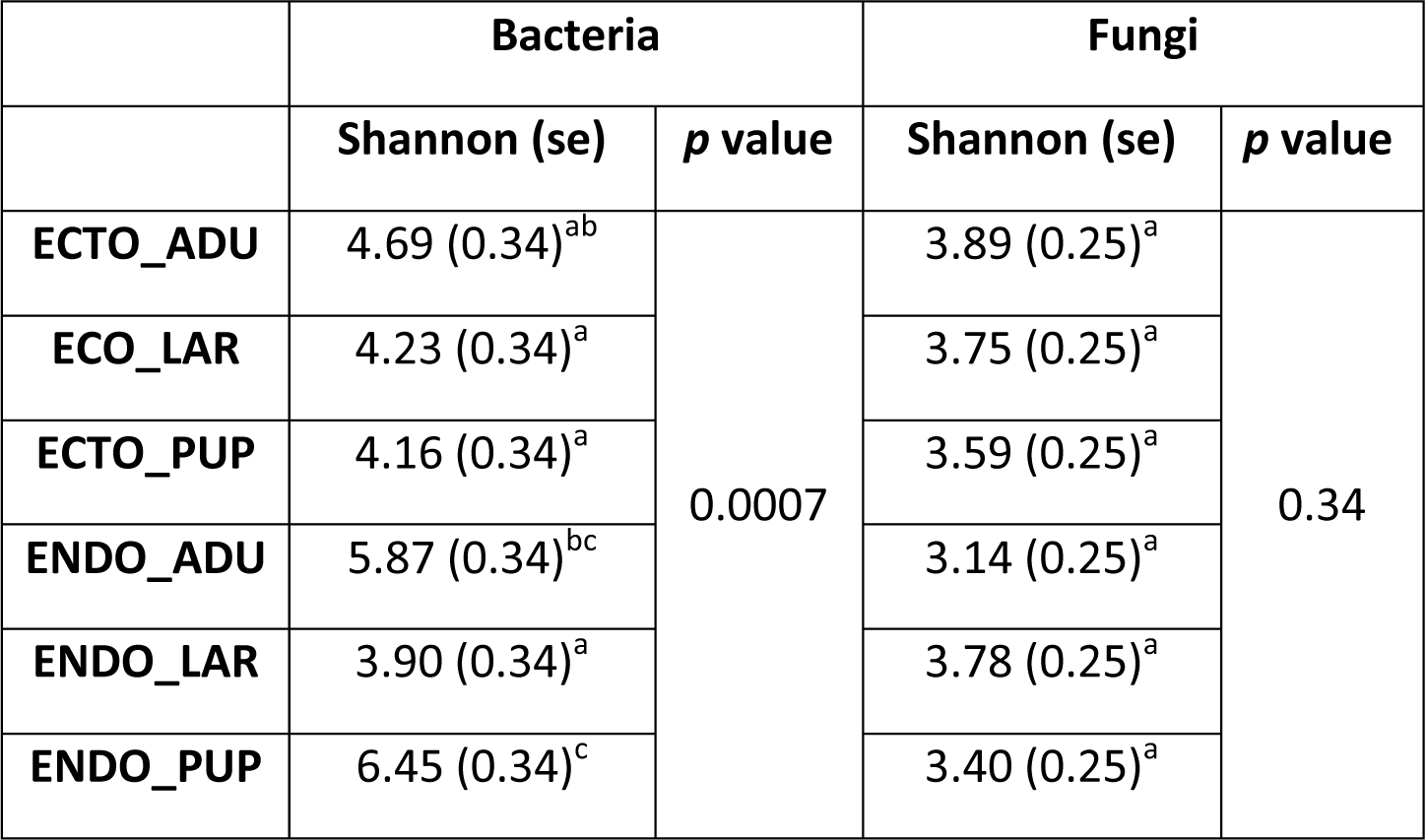
Shannon index for bacterial and fungal microbiota throughout *D. simplex* development. The Shannon index was calculated on OTUs obtained from equalized dataset. The mean of three replicates is presented in the table, with the standard error in parenthesis. ANOVA test was performed to identify significant differences between samples. ECTO = ectomicrobiota, ENDO = endomicrobiota, ADU = adults, LAR = Larvae, PUP = Pupae.

|  | Bacteria |  | Fungi |  |
| --- | --- | --- | --- | --- |
|  | Shannon (se) | *p* value | Shannon (se) | *p* value |
| ECTO_ADU | 4.69 (0.34)<sup>ab</sup> | 0.0007 | 3.89 (0.25)<sup>a</sup> | 0.34 |
| ECO_LAR | 4.23 (0.34)<sup>a</sup> |  | 3.75 (0.25)<sup>a</sup> |  |
| ECTO_PUP | 4.16 (0.34)<sup>a</sup> |  | 3.59 (0.25)<sup>a</sup> |  |
| ENDO_ADU | 5.87 (0.34)<sup>bc</sup> |  | 3.14 (0.25)<sup>a</sup> |  |
| ENDO_LAR | 3.90 (0.34)<sup>a</sup> |  | 3.78 (0.25)<sup>a</sup> |  |
| ENDO_PUP | 6.45 (0.34)<sup>c</sup> |  | 3.40 (0.25)<sup>a</sup> |  |

For the fungi, a total of 44,377 high-quality-filtered sequences were obtained after quality control and equalization, with an average read length of 447 bp. Clusterization at a 97% pairwise-identity threshold generated 1623 OTUs. No significant difference in Shannon’s diversity index is observed between fungal community structures of each sample (Table 1). After analysis, a suitable number of sequences were obtained to characterize the eastern larch beetle bacterial and fungal microbiota.

### Variation and taxonomical composition of the microbial communities across the eastern larch beetle developmental stages

Bacterial community differences within the beetle ontogeny are observed on the similarity dendrogram (Fig. 1). Samples associated with the endomicrobiota of the adults and pupae constitute a cluster significantly different from other samples. Additionally, the ectomicrobiota of the adults is creating a distinct cluster from the other ectomicrobiota samples. Accordingly, the ectomicrobiota of the pupae and larvae are grouped along with the endomicrobiota of the larvae, showing significant differences in the bacterial community for the interior of the insect at the larval stage.

**Figure 1.**
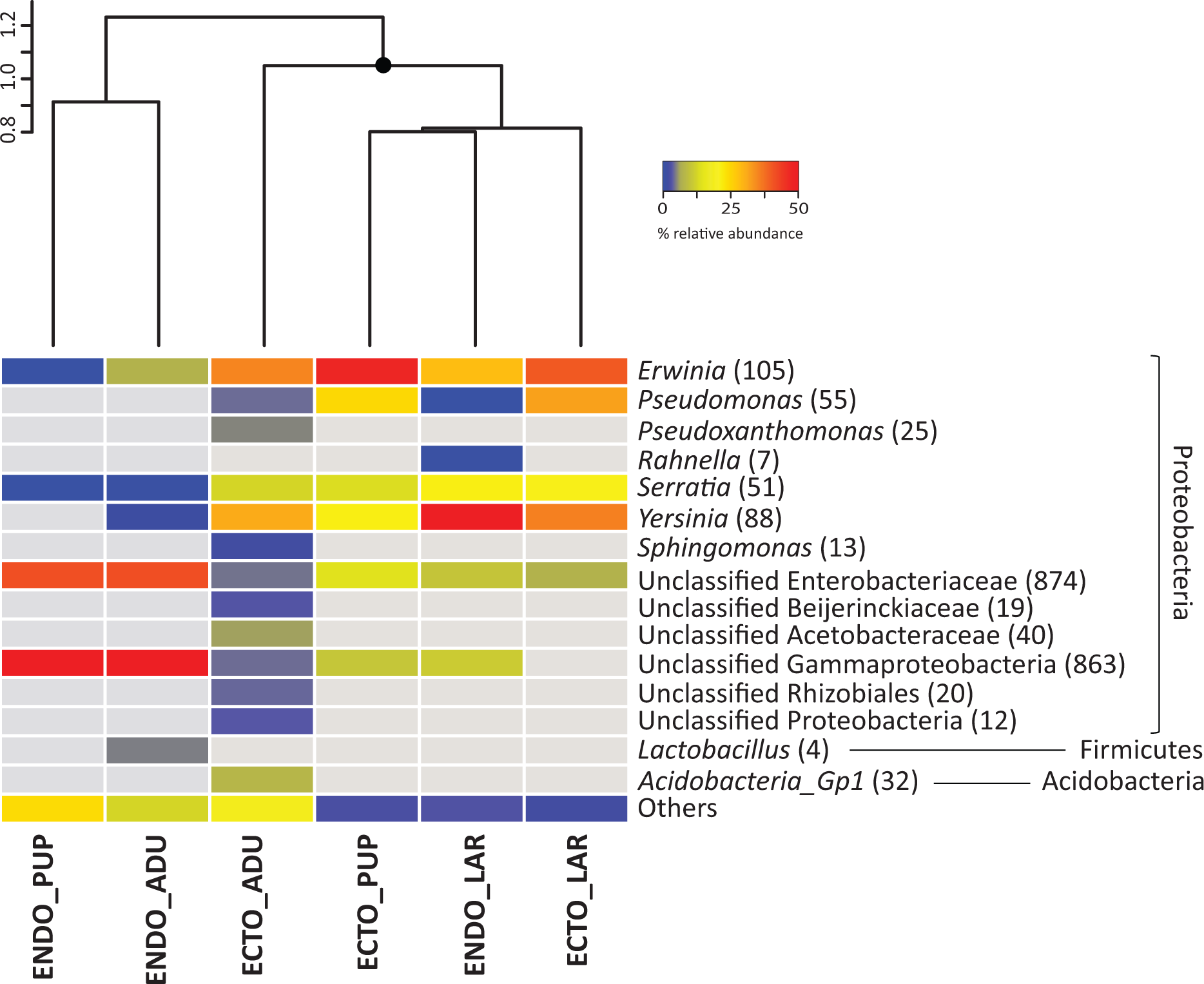
Bacterial community associated with different *D. simplex* developmental stages. Based on equalized dataset. The mean relative abundance of the three replicates is presented. Abundant OTUs (≥1% of relative abundance in one sample) are presented, non-abundant OTUs (<1%) are grouped in the category others. The grey color represents the absence of OTUs in the samples. The number in parenthesis represents the abundance of OTUs. Similarity cluster (UPGMA) grouping the different samples is presented above the heatmap, with significant (≥95%) node mark as black circles. ECTO = ectomicrobiota, ENDO= endomicrobiota, ADU = adults, LAR = Larvae, PUP = Pupae.

To identify the microbial composition of these communities, all identified bacterial OTUs were taxonomically assigned using the RDP classifier. Almost all bacterial OTUs were Proteobacteria, but some Firmicutes and Acidobacteria were also found. Among the Proteobacteria, Alphaproteobacteria and Gammaproteobacteria were represented. Figure 1 shows the taxonomical identification for the abundant bacterial OTUs (≥1% of the relative abundance per sample). For the ectomicrobiota, a few bacterial genera are abundant at all developmental stages, such as *Erwinia, Pseudomonas, Serratia, Yersinia* and unclassified OTUs belonging to the Enterobacteriaceae family. Altogether, these OTUs represent 60% of the total abundance in adults, and 98% and 90% for the larvae and pupae respectively. Additionally, unclassified OTUs belonging to the Gammaproteobacteria class were also identified in samples associated with the adults and pupae, with relative abundance up to 8%. The ectomicrobiota of the adults, which exhibited the highest specific richness of all developmental stages, included other lower abundance distinct OTUs: *Pseudoxanthomonas* and *Acidobacteria* Gp1, Alphaproteobacteria such as *Sphingomonas,* unclassified Beijerinckiaceae, Acetobacteraceae, and Rhizobiales. These OTUs are explaining the separate cluster set by adult ectomicrobiota over other samples.

Some of the bacteria found in the ectomicrobiota of the larvae were also associated with their endomicrobiota, such as *Erwinia, Serratia*, and *Yersinia,* representing 82% of the relative abundance. The *Erwinia* genus was also identified in the endomicrobiota of the adults, but in lower abundance (6% of the samples abundance). Additionally, OTUs related to the *Lactobacillus* genus were also identified in these samples with 4% of the total abundance. Several unclassified OTUs linked to Enterobacteriaceae and Gammaproteobacteria were documented in samples associated with the endomicrobiota of the adults and pupae, representing, respectively, 81% and 84% of relative abundance. Some of these OTUs were also represented in the endomicrobiota of the larvae, but in lower abundance (15%). Finally, other non-abundant OTUs were also identified throughout the beetle developmental stages.

The similarity dendrogram for fungal communities (Fig. 2) shows different grouping patterns than for the bacterial communities. Indeed, the pupae samples (ecto- and endomicrobiota) are arranged in a cluster significantly different from the other samples. Additionally, samples associated with the ectomicrobiota of the adults and larvae constitute a cluster significantly distinct from those of their endomicrobiota.

**Figure 2.**
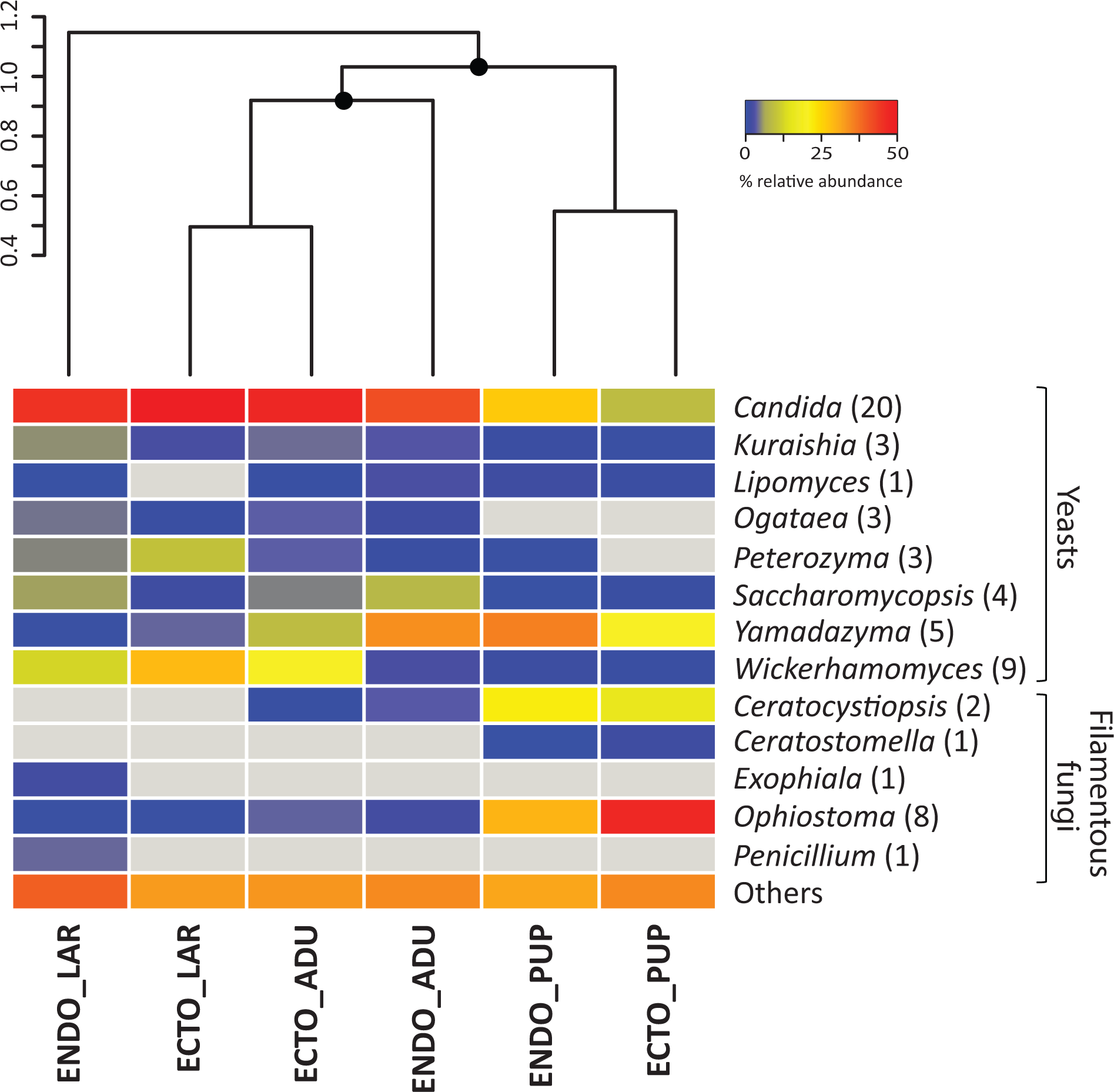
Fungal community related to different *D. simplex* developmental stages. Based on equalized dataset. The mean relative abundance of the three replicates is presented. Abundant OTUs are presented, non-abundant OTUs are grouped in the category others. The grey color represents the absence of OTUs in the samples. The number in parenthesis represents the abundance of OTUs. Similarity cluster (UPGMA) grouping the different samples is presented above the heatmap, with significant (≥95%) node mark as black circles. ECTO = ectomicrobiota, ENDO = endomicrobiota, ADU = adults, LAR = Larvae, PUP = Pupae.

All fungal OTUs were taxonomically identified by BLASTN against the NCBI database. All OTUs belong to the Ascomycota phylum. Among them, yeasts, all from the Saccharomycetes class, and filamentous fungi were identified. Figure 2 shows the taxonomical identification for the abundant fungal OTUs. Yeasts were predominant in samples associated with the adults and larvae, whereas filamentous fungi were mainly related to pupal samples, both for the ectomicrobiota and the endomicrobiota. *Candida* was the most abundant genus found associated with the adults and larvae ectomicrobiota (41% and 45% respectively), followed by *Wickerhamomyces* (13% and 19%). Others genera were also found, such as *Kuraishia, Ogataea, Peterozyma, Saccharomycopsis*, and *Yamadazyma*, with abundances ranging from 2% to 7%. The ectomicrobiota of pupae included only two abundant yeasts, *Candida* (7%) and *Yamadazyma* (14%), and two abundant filamentous fungi belonging to the Ophiostomatoid group, *Ceratocystiposis* (11%) and *Ophiostoma* (41%). Only *Ophiostoma* was associated in abundance with the adult ectomicrobiota samples.

Yeasts belonging to the *Candida* genus were also prevalent in the endomicrobiota at all developmental stages, with relative abundances ranging from 17% to 39%. Additionally, the *Yamadazyma* genus was also identified in samples from the adults and pupae (24% and 26%, respectively). The other yeasts genera identified in the ectomicrobiota of the adults and larvae were also present in their endomicrobiota (1% to 9%). Two main filamentous fungi identified in the ectomicrobiota of the pupae, *Ceratocystiopsis* and *Ophiostoma*, were also identified in abundance in their endomicrobiota samples, with, respectively, 14% and 19%. These filamentous fungi were also discovered in the endomicrobiota of the adults, but with lower abundances. As the adult’s samples, few filamentous fungi were associated with the larvae, identified as *Exophila* (2%) *and Penicillium* (3%).

A PERMANOVA analysis was performed to explain if the variability was triggered by both the developmental stages and the microbiota location. For bacteria and fungi, the developmental stage explained a higher percentage of the variation (43% (*p* < 0.001) for bacteria and 59% (*p* < 0.001) for the fungi) than the microflora (14% (*p* < 0.001) for bacteria and 9% (*p* < 0.006) for the fungi).

## Discussion

Bark beetles form associations with bacteria, filamentous fungi, and yeasts. These microorganisms play a major role in the beetle development and help the insect colonize the subcortical environment of the host tree (Six, 2013; Hofstetter *et al.*, 2015). They can be acquired either from the environment or directly from their parents by vertical transmission (Gibson & Hunter, 2010). To better understand the symbiotic relationship between a beetle and its associated microorganisms, it is important to acquire a comprehensive portrait of the microbiota associated with the insect. Accordingly, the beetle microbiota, or at least the relative abundance of its various members, should change over the ontogeny of the insect in function of beetle requirements, resulting in a successional microbial assemblage. In this context, we characterized the microbiota associated with the developmental stages of the eastern larch beetle by high throughput sequencing to observe the progression of the microbiota over time. Because the surface and interior of the insect represent two distinct microbial communities, we investigated both populations separately.

As expected, both similarity dendrograms and taxonomical identifications revealed a succession of the abundant microorganisms associated with the eastern larch beetle throughout its ontogeny. Accordingly, the developmental stages explain a high proportion of the observed variations, as the PERMANOVA analyses show. These differences are probably related to the beetle requirements over the developmental process. Figure 3 shows the proposed succession of abundant microorganisms and their possible functions according to the developmental stages and the microbiota of *D. simplex.* Non-abundant OTUs were still present in the samples, but not represented in the schema. This diagram is based on the sequencing results obtained and existing literature on beetle-associated microorganisms. Attack of tree and development of the eastern larch beetle start with a dispersion period. After the selection of a suitable host, pioneer beetles bore holes in the tree trunk and build-up galleries in the phloem layer (Langor & Raske, 1987a). To successfully attack and colonize the subcortical environment of the host tree, the insects need to overcome the tree defense system, for instance by clogging resin ducts and degrading terpenoid compounds (Paine *et al.*, 1997; Adams *et al.*, 2011). The microbiota is essential in these defensive actions (Six & Wingfield, 2011; Six, 2013). Accordingly, pioneer beetles need to carry, mainly under their elytra, their associated microorganisms playing a key role in the protection from tree defense mechanisms (Durand *et al.*, 2015). Ophiostomatoid fungi have been frequently reported for their ability to colonize the resin ducts of conifer trees, enabling the insect to settle in the phloem without being trapped in the resin (Paine *et al.*, 1997; Lieutier *et al.*, 2009; Six & Wingfield, 2011). Indeed, the *Ophiostoma* genus was the only filamentous fungi identified in abundance (3%) in the ectomicrobiota of the adults. The beetle at this stage would transport and promote this filamentous fungus on its exoskeleton to weaken the tree defense system and thus benefit from it by successfully colonizing the host tree. Additionally, *Pseudomonas* and *Serratia*, which were reported to play a role in the degradation of terpenoid compounds in other *Dendroctonus* species, were identified in high proportion in the ectomicrobiota of *D. simplex* adults (Morales-Jimenez *et al.*, 2012; Adams *et al.*, 2013; Boone *et al.*, 2013). Yeasts, such as *Ogataea pini*, also abundant in the ectomicrobiota of adults, are able to tolerate and grow in the presence of some terpenoid compounds, but no direct evidence of degradation has been demonstrated (Davis & Hofstetter, 2011). Indeed, the beetle could be associated with these bacteria and yeast to benefit from the detoxification of the environment. With this assemblage of microorganisms carried on its cuticle, the insect could face the tree defense system and successfully colonize the phloem. In return, microorganisms get access to a new niche.

**Figure 3.**
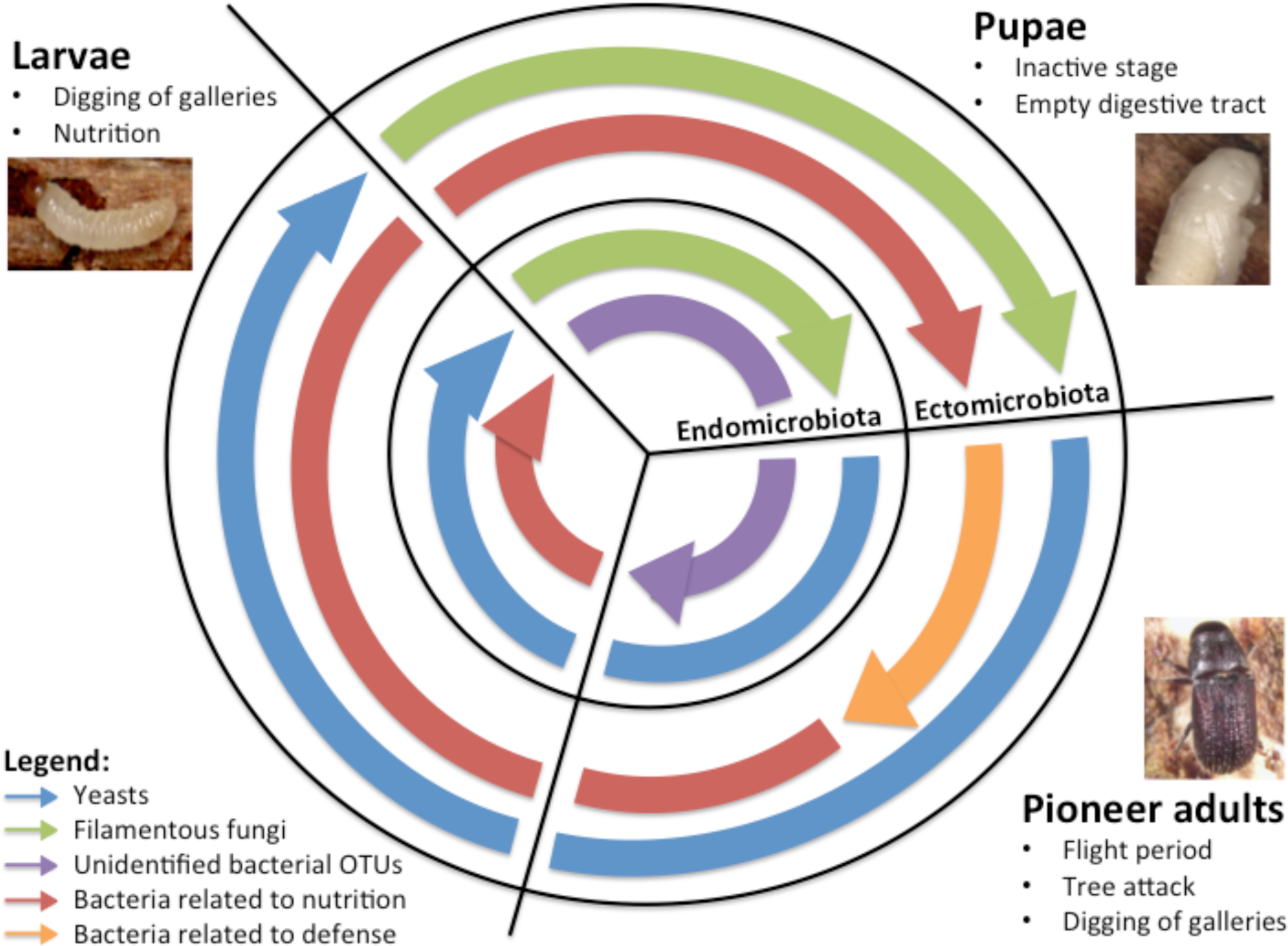
Overview of the proposed successional microbiota associated with *D. simplex*developmental stages and microflora, based on obtained results and literature. The predominant microorganisms for each developmental stage and microbiota are presented in the figure.

Apart from the plant protection mechanisms, insects are also confronted to antagonistic microorganisms under the bark, such as fungi. *Pseudomonas* and *Serratia* genera have been reported to express antifungal activities against antagonistic fungi and are believed to play a major role in the beetle defense against pathogenic microorganisms under the bark (Cardoza *et al.*, 2006; Winder *et al.*, 2010). Moreover, yeasts and filamentous fungi associated with *Dendroctonus* species are also able to produce volatile compounds that modulate the growth of other fungi under the bark (Adams *et al.*, 2008; Cale *et al.*, 2016). *Ogataea pini*, associated with a few *Dendroctonus* species, produce volatile compounds that inhibit the growth of the entomopathogenic fungus *Beauveria bassiana* (Davis & Hofstetter, 2011). Accordingly, the *Ogataea* genus was identified in abundance in the ectomicrobiota of the adults. Based on the published functions and the presence of many microorganisms associated with the ectomicrobiota of the adults, we hypothesize that *D. simplex* is carrying on his exoskeleton a microbial assemblage implicated in the defense mechanisms of the insects, protecting them during the colonization.

Numerous studies report the implication of microorganisms in beetle nutrition, completing the nutrient-poor phloem diet (Paine *et al.*, 1997; Adams & Six, 2007; Six & Wingfield, 2011; Popa *et al.*, 2012; Davis, 2014). As beetle pioneers colonize a new environment, they should bring along these microorganisms on their exoskeleton to colonize the galleries they are digging. They can then proliferate under the bark, enabling the next generation of insects to feed on them. Such microbial gardening has been observed in ants, termites, and ambrosia beetle (Mueller & Gerardo, 2002; Mueller *et al.*, 2005). Many authors hypothesize that filamentous fungi could be a nutritional source for beetles, mainly by providing nitrogen and sterols, essential for oogenesis and larval development (Klepzig & Six, 2004; Bentz & Six, 2006; Adams & Six, 2007). Moreover, *Candida* species isolated from Coleoptera degrade many compounds, such as sugar and cellulose, and produce vitamins (Chararas *et al.*, 1983). More recently, the implication of associated yeasts in beetle nutrition has been hypothesized (Davis, 2014; Durand *et al.*, 2017). Accordingly, yeasts such as *Candida* were found in abundance in feeding developmental stages (adults and larvae) while being underrepresented in the inactive stage (pupae). This finding supports the implication of yeasts in beetle nutrition. Bacteria are also essential for the beetle diet, acting mainly as nitrogen fixators and cellulose degraders (Popa *et al.*, 2012). *Rahnella aquatilis*, associated with many *Dendroctonus* species, can fix nitrogen, leading authors to hypothesize that this bacterium could concentrate nitrogen to fulfil insect needs (Morales-Jimenez *et al.*, 2009; Morales-Jimenez *et al.*, 2012). Additionally, many *Pseudomonas* species, such as *Pseudomonas fluorescens*, also isolated from *Dendroctonus* species, have cellulolytic activity (Morales-Jimenez *et al.*, 2012; Hu *et al.*, 2014; Briones-Roblero *et al.*, 2017). Bacteria belonging to this genus could be implicated in cellulose breakdown, helping the insect to assimilate recalcitrant carbon sources. Vitamins and amino acids are also thought to be supplemented by bacteria, but no direct evidence is yet available (Gibson & Hunter, 2010). We hypothesize that filamentous fungi, yeasts, and bacteria providing nutritional benefits are transported by pioneer adults of *D. simplex*, and are inoculated in the galleries during their construction, so the next generation can feed on them to complete their development, explaining their predominant presence in the ectomicrobiota of adults. As demonstrated before, the beetle’s ectomicrobiota enclosed specific OTUs, distinct from the galleries samples (Durand *et al.*, 2015). As pioneer beetles attack and colonize new trees, a selection of beneficial microorganisms by the insect could occur, leading them to a successful attack.

Other bacteria were identified in adults’ ectomicrobiota, such as *Erwinia, Pseudoxanthomonas, Yersinia*, and some unclassified bacteria, but no specific function is yet attributed to them. Some authors suggested that they could be involved in beetle defense or nutrition, or even play some role in the pheromone synthesis or the mediation of interactions among the microbiota (Gibson & Hunter, 2010; Popa *et al.*, 2012; Hofstetter *et al.*, 2015). Additionally, these bacteria could also be associated with the beetle’s galleries rather than transported by the insect, and colonize the ectomicrobiota after galleries construction, as some of them were previously identified in *D. simplex* galleries (Durand *et al.*, 2015).

After reproduction and hatching of eggs, larvae construct their horizontal galleries and feed on phloem throughout their development (Langor & Raske, 1987b). Accordingly, microorganisms identified in high proportion in the larval stage should shift and be related to nutritional benefits, both for the ectomicrobiota, as they colonize the galleries, and the endomicrobiota, as they feed on them. Indeed, yeasts were predominant in both microbiota. Furthermore, the same yeast genera were also found in the adults and larvae, with different abundance for the endomicrobiota, likely reflecting different nutritional requirements for these two developmental stages. Moreover, bacteria related to nutrition identified in adults were also present in the ecto- and endomicrobiota of larvae, supporting the hypothesis of their nutritional benefits. Indeed, results for larvae microbiota support the hypothesis that pioneers transport microorganisms on their exoskeleton to enable them to grow in the galleries and the next generation to feed on them. The Shannon diversity index is also lower in the larvae endomicrobiota compared to other developmental stages. This finding suggests that a fewer number of bacterial species is required for the larval nutrition when the insect evolves under the bark, as they live a simpler life once the attack and colonisation of the tree is achieved.

In preparation for the transition to the pupal stage, a larvae excavates a pupal chamber, stops feeding, empty his gut and blocks the entrance hole with frass (Langor & Raske, 1987b). Accordingly, filamentous fungi were predominantly associated with this stage, for both the ecto- and endomicrobiota, whereas yeasts were underrepresented. This further supports a model where yeasts play a fundamental role in nutrition. The pupa is inactive under the bark, and so this developmental stage is more vulnerable. Pupal chambers harbor spores of filamentous fungi, and it was hypothesized that beetles could acquire them at this stage (Six, 2003). Associated filamentous fungi produce antifungal compounds that inhibit the growth of antagonistic fungi, protecting the beetle under the bark (Paine *et al.*, 1997; Cale *et al.*, 2016). Pupae may form an association with these filamentous fungi for their protection during this crucial period of development, explaining the presence of more genera of filamentous fungi associated with this developmental stage. The change from active to passive behaviour could then explain the significant modification in fungal community structure of *D. simplex* pupae. Additionally, the bacterial population exhibits a similar shift in its abundant microorganisms, as bacteria involved in nutritional functions were underrepresented. Unclassified Enterobacteriaceae and Gammaproteobacteria were mainly associated with pupae. Unfortunately, no particular function has been attributed to these bacteria yet. Our results suggest that there are probably endosymbiotic bacteria as they are found in pupae after the beetle empties its gut, and also in pioneer beetles. Additionally, these unclassified groups were recovered from the endomicrobiota of *D. simplex* in our previous study on hybrid larches (Durand *et al.*, 2015). These OTUs appear to represent novel species, as they do not correspond to known bacteria in public databases.

After the pupal stage, the insect transforms into adult and hibernates in the pupal chamber for winter. Before the flight period in the spring, the new generation of insects needs to feed to reach its sexual maturity (Langor & Raske, 1987b). Apart from the phloem, insects may feed on filamentous fungi found in their pupal chamber, as they were reported to contain a high concentration of sterols, but mainly on yeasts as they are found in abundance in the adult’s endomicrobiota and in the galleries (Bentz & Six, 2006; Adams & Six, 2007; Durand *et al.*, 2017). Accordingly, mainly the unclassified Enterobacteriaceae and Gammaproteobacteria were identified in these samples, and the bacteria related to nutrition were underrepresented, also supporting the hypothesis of a feeding function for the yeasts. The next generation will then emerge from the bark and attack a new tree.

When considering the whole developmental process and the microorganisms altogether, we can hypothesize that bark beetle pioneers carry the microbial assemblage needed for a successful attack and the complete development cycle occurring under the bark. Indeed, all microorganisms with possible functions were identified in the adult samples. According to the beetle’s requirements, the abundance of microorganisms shifts between the developmental stages, as their functions differ. Some of these microorganisms could be acquired by the new generation by vertical transfer or could be acquired by horizontal transfer as they colonize the galleries.

Investigating the structure of the microbial community (bacteria, filamentous fungi, and yeasts) associated with the eastern larch beetle revealed significant changes in their relative abundance over the various stages. Changes in bacterial and fungal community structures seem to be triggered by the physiological origin (ecto and endo) and by the developmental stages of the insect. These variations can be explained by the succession of functions required by each developmental stage, and beetle requirements. These results suggest that a symbiotic association between the eastern larch beetle and some of these microorganisms take place and that this *D. simplex* symbiotic complex is helping the insect to colonize its host tree and survive under subcortical conditions.

## Acknowledgments

We would like to thank Amélie Bergeron for her help with samples preparation and DNA extraction. We would also like to thank Fabrice Jean-Pierre, Jean-Philippe Buffet and Marie-Christine Groleau for their technical assistance.

